# Efficient Immune Cell Genome Engineering with Improved CRISPR Editing Tools

**DOI:** 10.1101/2020.02.13.947002

**Authors:** Waipan Chan, Rachel A. Gottschalk, Yikun Yao, Joel L. Pomerantz, Ronald N. Germain

## Abstract

CRISPR (clustered regularly interspaced short palindromic repeats)-based methods have revolutionized genome engineering and the study of gene-phenotype relationships. However, modifying cells of the innate immune system, especially macrophages, has been challenging because of cell pathology and low targeting efficiency resulting from nucleic acid activation of sensitive intracellular sensors. Likewise, lymphocytes of the adaptive immune system are largely refractory to CRISPR-enhanced homology-directed repair (HDR) due to inefficient or toxic delivery of donor templates via transient transfection methods. To overcome these challenges and limitations, we developed three improved methods for CRISPR-based genome editing using a hit-and-run transient expression strategy to minimize off-target effects and generate more precise genome editing. Overall, our enhanced CRISPR tools and strategies designed to tackle both murine and human immune cell genome engineering are expected to be widely applicable not only in hematopoietic cells but also other mammalian cell types of interest.

All animal experiments were done in accordance with the guidelines of the NIAID/NIH Institutional Animal Care and Use Committee.

## Introduction

The bacterial innate defense system CRISPR was discovered more than 3 decades ago (1), yet this game-changing genome engineering technology was not harnessed for human gene editing applications until a series of landmark studies published in 2013 (2–4). Since then, there has been an exponential increase of CRISPR-related publications, mainly focusing on the use of the Cas9 nuclease that comes with unpredictable off-target effects (5–7), hindering its direct application to the clinic. Many research groups have developed novel methods to minimize these potential side-effects (8–14). The Cas9 nickase method is a promising approach due to its strict pairing distance requirement, which makes off-target pairing very unlikely to occur and therefore minimizes off-target effects. Another straight-forward strategy to reduce CRISPR-Cas9 off-target effects is to limit the expression duration of Cas9 or guide-RNA (gRNA), the same rationale behind the use of Cas9 inhibitors. While this can be achieved using transient transfection methods, the on-target editing efficiency can be compromised (15) as compared to the stable lentiviral transduction methods.

Despite many advances in CRISPR technology, to date there have been very few reports of using the Cas9 nickase strategy for immune cell (myeloid cell or lymphocyte) genome engineering. Macrophages are a major cell type involved in innate immune responses and have been especially difficult to modify using available CRISPR methods. In particular, genome editing of macrophage cell lines using all-in-one CRISPR-Cas9 techniques has been inefficient (16) and although newer methods have yielded improved results (17–19), some of these approaches require antibiotic selection and clonal isolation that dramatically delays the phenotyping timeline and risks the loss of original cell properties. To enable high-throughput CRISPR-Cas9 genome-wide knockout (KO) library screens, the murine RAW264.7 macrophage (RAW) cell line has been modified to constitutively express Cas9 for efficient genome editing (20). While constitutive Cas9 and gRNA expression can lead to the accumulation of off-target mutations in the long run, the doxycycline-inducible system pCW-Cas9 (21) helps resolve both issues by controlling the genome editing time window to minimize long-term off-target effects and allowing immediate phenotyping before the onset of potential compensating mechanisms. However, the lack of a live cell marker makes it difficult to identify and isolate the Cas9-expressing population for further analysis. Here we describe a modified method to enable doxycycline-induced Cas9 expression to be tracked by EGFP fluorescence in live single cells and utilize this system to effectively engineer the RAW cell line.

Macrophages are also problematic to engineer due to their innate immune sensors. Lentiviral particles produced from packaging cells such as the HEK293T cell line are usually filtered or spun to remove cell debris. However, this process does not remove many small molecules and exosomes secreted from the exhausted and dying packaging cells. Lentiviral supernatant containing these uncharacterized materials are often directly applied to cell culture and the potential side effects of the treatment due to sensor signaling are often ignored. Here we improve the purity of lentiviral particles to enhance macrophage cell viability post-transduction. With an optimized protocol to transduce primary bone marrow derived macrophages (BMDM) derived from the Cas9 mice (22), we validated our cell-line RAW cell CRISPR-KO phenotypes using the same set of targeting gRNAs with primary macrophages.

In contrast to macrophages, lentiviral-mediated CRISPR-KO is highly efficient in human T lymphocytes (20,21). However, this approach is not suitable for CRISPR-HDR applications due to the random integration of the HDR donor template that can create undesired genomic instability. Transient transfection delivery methods such as lipid-based reagents and electroporation systems either yield low transfection efficiency or a high percentage of cell death. Furthermore, compared to the high targeting efficiency (80∼90%) of CRISPR-induced NHEJ (non-homologous end joining), the targeting efficiency of CRISPR-induced HDR is generally inefficient (10∼20% for large fragment insertion) (23). Many reported HDR applications relevant to immune cell manipulation involve targeting embryos (24) or newborn mice (25), not mature immune T cells. To achieve efficient and precise genome modifications in human Jurkat T cells as a model for mature lymphocyte engineering, we adopted the Cas9 nickase strategy and describe here an antibiotic selection strategy that helps enrich specifically for rare HDR clones following plasmid co-transfection. In addition, we further improved this method to minimize the genomic footprint created by the antibiotic resistance marker and make it possible for future rounds of genome editing in the same cells using the same strategy.

Another issue is enhancement of the efficiency of directed templated mutagenesis. While the NHEJ inhibitor SCR7 (26) and the HDR stimulatory compound RS-1 (27) were reported to enhance CRISPR-Cas9-induced HDR efficiency, the key to shifting the balance between the NHEJ and the HDR DNA repair mechanisms in favor of the latter outcome seems to rely on a high molar ratio of donor DNA to Cas9-gRNA ribonucleoprotein (RNP). This was demonstrated in human primary T cell engineering using 1e6 MOI of AAV6 (adeno-associated virus type 6) transduction to deliver a high copy number of HDR donor templates per target cell (28). AAV is a small single-stranded DNA virus that infrequently integrates into the host genome, making it a safe delivery platform for the HDR donor template. However, the production and purification process of AAV particles is relatively labor-intensive and often requires industrial assistance, which significantly increases the cost and reduces the flexibility of preliminary experiments in the laboratory setting. In addition, the very limited payload of the AAV vector makes it impossible to fit Cas9, gRNA expression and donor template in one AAV backbone. In comparison, lentiviral particles are easy to produce and purify in most laboratories with relatively low cost. Furthermore, the payload of lentiviral particles is large enough to carry Cas9, gRNA expression and donor template all-in-one. Despite these advantages, random integration of lentiviral DNA into the host genome makes it problematic as a platform to deliver the HDR donor template. Here we overcome this limitation by the use of integrase-deficient lentiviral particles (29) and report two examples of this novel approach in human Jurkat T cells.

Finally, all these methods involve either gene inactivation or modification. There are numerous situations in which augmented expression of a given gene would be desirable in an experimental setting. While global methods for gene activation using a so-called CRISPRa strategy have been reported (30–32), efficient induction of selective gene expression is not yet a commonly used method nor do simple tools exist for this purpose. Here we describe a lentiviral-based system for individual gene activation control and show its utility as applied to the human Jurkat T cell model. In combination with the previously noted improvements for macrophage genome editing, these various new vectors and strategies constitute an effective suite of methods for the manipulation of both innate (macrophage) and adaptive (lymphocyte) immune cells that offer benefits when applied to a wide variety of cells types.

## Materials and Methods

### Gibson Assembly cloning of donor and lentiviral expression plasmids

The pCW-Cas9-2A-EGFP plasmid was constructed by replacing the FseI-BamHI fragment in the pCW-Cas9 plasmid (Addgene plasmid #50661) (21) with the T2A-EGFP fragment. The LCv2B plasmid was constructed by replacing the BamHI-MluI fragment in the lentiCRISPRv2 plasmid (Addgene plasmid #52961) (33) with the P2A-BSD fragment. The LGB and the LGR plasmids were constructed by replacing the XbaI-Cas9-BamHI fragment in the LCv2B plasmid with the TagBFP and the TagRFP-T fragment respectively. The STAT3 HDR donor plasmid was constructed by replacing the NotI-XhoI fragment in the pcDNA3.1 plasmid with the indicated fragments. The BCL10 HDR donor plasmid was assembled based on a minimal PciI-BglII pcDNA3 fragment that contains only the ColE1 origin and the ampicillin resistance expression cassette. To construct the LCv2B-HDR-mCherry-2A-hCD3E plasmid, LCv2B plasmid containing either the CD3E gRNA-1 or 2 was linearized at the KpnI site upstream of the U6 promoter before the assembly of the donor template fragments. The LCv2S plasmid (Figure 5 - figure supplement 1A) was constructed by replacing the BamHI-MluI PuroR fragment in the lentiCRISPRv2 plasmid with a 24-bp bridging fragment via sticky-end ligation. To construct the LCv2S-HDR-tBFP-hRelA plasmid, LCv2S plasmid containing the RelA gRNA was linearized at the same KpnI site to allow for the assembly of the donor template fragments. The LCv2R plasmid was assembled by adding the P2A-mScarlet-I fragment to the BamHI-linearized LCv2S backbone. The LCv2E plasmid was assembled using In-Fusion cloning by adding the P2A-tEGFR fragment to the BamHI-linearized LCv2S backbone. The LES2A-CRE plasmid was assembled by replacing the XbaI-EcoRI fragment in the lentiCas9-Blast plasmid (Addgene plasmid #52962) (33) with the mScarlet-I-P2A-CRE fragment. The Lenti-CMV-mCherry-P2A-CRE (aka pLM-CMV-R-Cre) plasmid was a gift from Michel Sadelain (Addgene plasmid #27546) (34). The LentiSAMPHv2 plasmid was assembled by adding the P2A-MS2-p65-HSF1 fragment amplified from the lentiMPH v2 plasmid (Addgene plasmid #89308) (32) to the BsrGI-linearized lentiSAMv2 backbone (Addgene plasmid #75112) (32). All constructs described in this paper will be deposited with Addgene and become available to qualified investigators.

### CRISPR gRNA target design and cloning

All gRNAs were designed based on the analysis results from the Zhang Lab online platform (crispr.mit.edu). For the DUSP gene knockout application, we selected the gRNA with the best score that targets downstream of the start codon of each DUSP gene. For the STAT3 HDR application, we selected a gRNA pair with the best score that targets near the start codon. For the BCL10 HDR application, we selected top five scoring gRNA pairs that target near the start codon. In addition, we selected a gRNA pair (D3) that confers a small offset distance of 3 bp. Both the CD3E gRNA-1 and the RelA gRNA for the NIL delivery applications were designed to overlap with the start codon in each case in order to avoid further editing following the HDR event. With the same rationale in the case of CD3E gRNA-2, a PAM mutation was introduced into the LCv2B-HDR-mCherry-2A-hCD3E plasmid. The RelA gRNA A1 was adopted from the human GeCKOv2 library A (33). The PX461 and PX462 plasmids were gifts from Feng Zhang (Addgene plasmid #48140 & #48141) (35). The pKLV-BFP plasmid was a gift from Kosuke Yusa (Addgene plasmid #50946) (36). All lentiviral and other gRNA expression plasmids were prepared by sticky-end ligation before any further modification described. The PX462-hBCL10-gRNA-A1 plasmid was constructed by Gibson Assembly in the PciI-linearized PX461 backbone, followed by sub-cloning of the double gRNA expression fragment into the PX462 backbone with PvuI-XbaI sticky-end ligation. CRISPRa gRNAs were adopted from the Human CRISPR Activation Library (SAM – 2 plasmid system) from Feng Zhang (32).

### Production, purification and concentration of lentiviral particles

Lentiviral particles were produced in HEK293T cells following transient transfection of the packaging plasmids pVSVg (Addgene plasmid #8454) and psPAX2 (Addgene plasmid #12260), and the lentiviral expression plasmid in a 1:10:10 ratio using Lipofectamine 3000. Lentiviral supernatant was harvested 44 to 52 h post-transfection and spun at 500 X *g* for 5 minutes to clear cell debris. NIL particles were produced the same way except with the use of the psPAX2-D64V plasmid (Addgene plasmid #63586) (29) instead of the psPAX2 plasmid. All lentiviral particles other than those used in the RAW cell transductions and Figure 2B (top panel) were further purified and concentrated with the Lenti-X Concentrator reagent (Takara) according to the manufacturer’s instructions.

### Cell culture, transfection and antibiotic selection conditions

HEK293T, RAW264.7 and Jurkat cell lines were purchased from ATCC. Cell culture was performed in a humidified environment with 5% CO_2_ at 37°C. Both HEK293T and RAW264.7 cell lines were cultured in DMEM supplemented with 10% FBS, glutamine and 18 mM HEPES buffer. Jurkat cell lines were cultured in RPMI supplemented with 10% FBS, penicillin/streptomycin, glutamine and 50 μM β-mercaptoethanol (complete RPMI). For plasmid transfections of HEK293T and Jurkat cells, Lipofectamine 3000 (ThermoFisher) was used according to manufacturer’s protocol. Puromycin (0.5 to 3 μg/mL) or blasticidin (1 to 3 μg/mL) was added to cell culture medium 1 to 2 days after transduction of both RAW and Jurkat cell lines.

### Genomic DNA extraction, T7EN assay and sequencing

Genomic DNA was extracted from RAW, HEK293T or Jurkat cells using the QIAmp DNA Mini Kit (Qiagen) following the manufacturer’s protocol. Genomic regions of interest were then amplified by PCR for follow-up analysis. For the T7EN assay, equal amounts of genomic PCR products were used within each comparison group for the hybridization reaction in a thermocycler programmed to 95°C for 10 minutes, -2°C per second ramp to 85°C, -0.1°C per second ramp to 25°C, and then held at 4°C until a subsequent T7 Endonuclease I (NEB) treatment at 37°C for 30 to 90 minutes. 25 mM final concentration of EDTA was added to each reaction before agarose gel electrophoresis analysis. Indel percentage was determined as described previously (8). For the genomic DNA sequencing, each DUSP gene locus surrounding the CRISPR gRNA target site (approximately -500 bp to +500 bp) was amplified and cloned into a XhoI-NotI linearized pcDNA3.1 backbone by Gibson Assembly. Sequencing was performed with the BGH-R (Table 1) primer on 20 clones per gene.

**Table 1.**
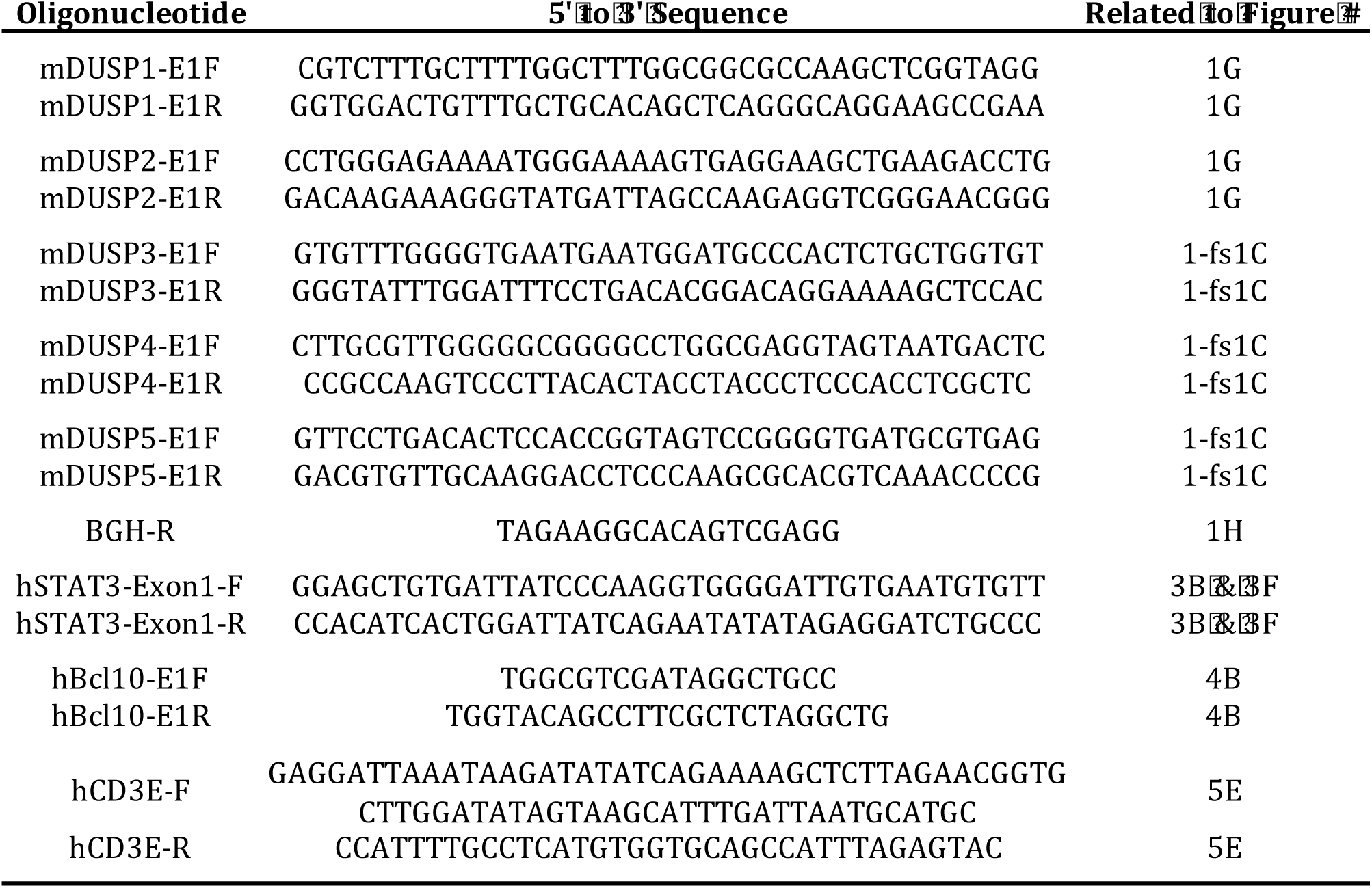
Summary of oligonucleotide sequences.

**Table 2.**
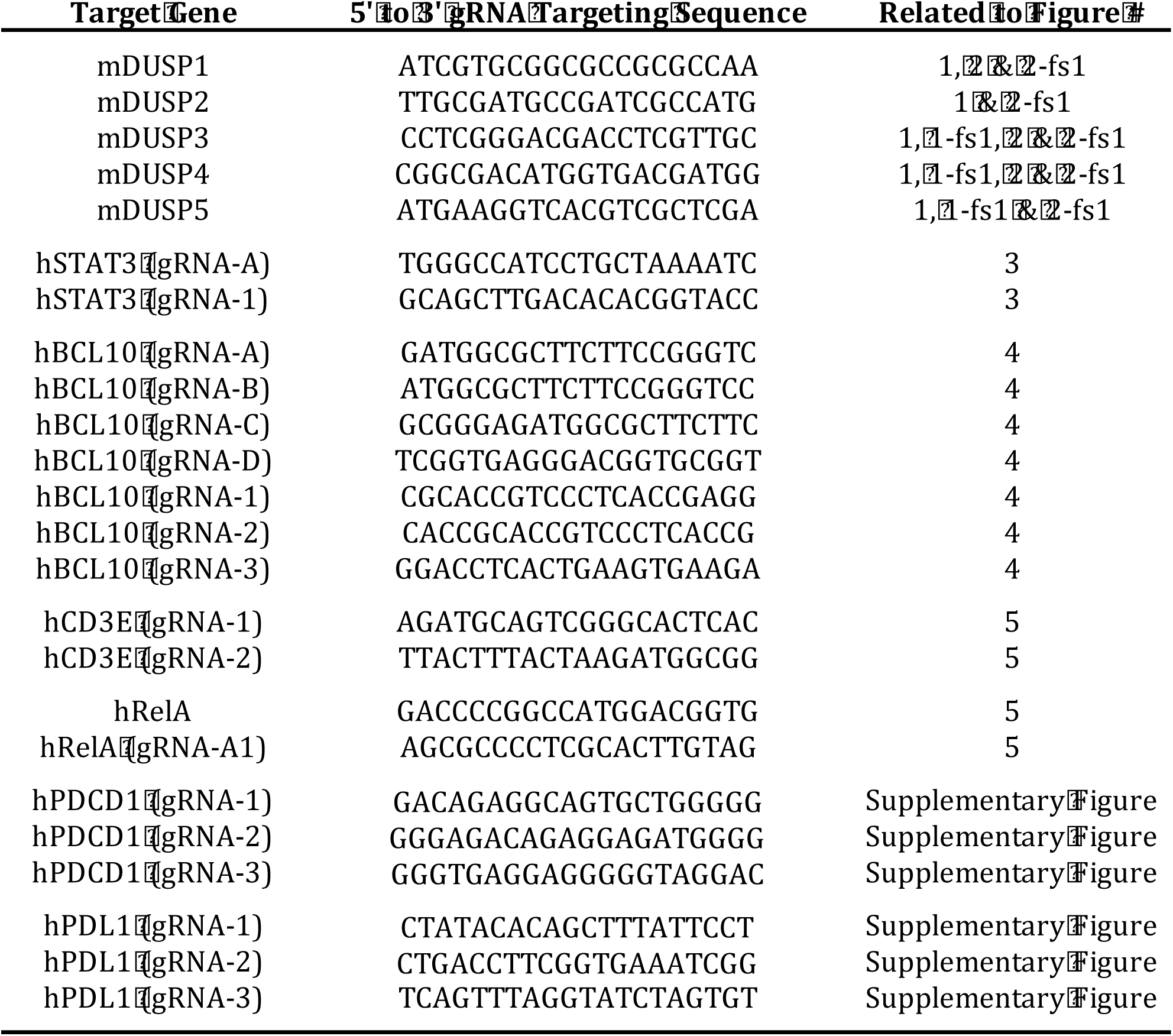
Summary of gRNA targeting sequences.

### Bone marrow isolation and intracellular Ki67 staining

Mice were maintained in specific-pathogen-free conditions and all procedures were approved by the NIAID Animal Care and Use Committee (National Institutes of Health, Bethesda, MD, ASP LISB-4E). Bone marrow progenitors isolated from C57BL/6 mice (Jackson Laboratories) and Rosa26-Cas9 B6J mice (Jackson Laboratories #026179) were differentiated into BMDM during a 7-day culture in complete Dulbecco’s modified Eagle’s medium (10% FBS, 100 U/mL penicillin, 100 U/mL streptomycin, 2 mM L-Glutamine, 20 mM HEPES) supplemented with 60 ng/mL recombinant mouse M-CSF (R&D systems). For analysis of proliferation, cells were harvested at the indicated day of the BMDM culture, were processed using BD Cytofix/Cytoperm reagents as directed, blocked and stained with anti-Ki-67 (BD Pharmingen).

### Macrophage TLR4 activation and intracellular phospho-p38 staining

BMDM or RAW cells were stimulated with TLR4 ligand Kdo2-Lipid A (KLA, Avanti Polar Lipids). Ligand addition was staggered so that for all time points, cells were fixed at the same time by addition of paraformaldehyde to cell cultures at a final concentration of 1.6% (10 minutes). After one wash with PBS 1% FBS, cells were gently harvested from plates by scraping, were permeabilized overnight using ice cold MeOH at -20°C, blocked using 5% goat serum and Fc receptor specific antibody, and stained for 1 h at room temperature with anti-p38 (phospho-Thr180/Try182, BD 36/p38) antibody.

### Cell sorting, FACS analysis and live cell confocal imaging

Cell sorting was conducted by the Flow Cytometry Section, Research Technologies Branch of NIAID. FACS analysis data were collected using a BD LSRII or BD Fortessa Cell Analyzer and further processed with FlowJo. Human PDCD1 or PDL1 cell surface protein staining was analyzed using anti-PD-1 (ThermoFisher clone MIH4) or anti-PD-L1 (ThermoFisher clone MIH1) antibody respectively. Live cell confocal time-lapse imaging data were collected using a Leica SP8 microscope with a 63x NA 1.4 oil objective (Biological Imaging Section, Research Technologies Branch of NIAID). The imaging chamber (ThermoFisher 155411) was coated with poly-D-lysine (Sigma P7280) for 1 h at 37°C and washed twice with PBS. Cells were imaged in a heated 37°C environment with 5% CO_2_.

Imaging data were processed by Imaris (Bitplane).

### Chemical reagents, lysis buffer and western blots

Doxycycline hydrochloride (Sigma D3447) was reconstituted in DMSO at 50 mg/mL and stored at -20°C. Jurkat T cells were stimulated with 50 ng/mL PMA (Santa Cruz Biotechnology) and 1 μM ionomycin (Sigma) (37). RAW cells were lysed in a buffer containing 50 mM Tris pH7.5, 150 mM NaCl, 1 mM EDTA, 10% glycerol and 1% Igepal. Western blots were analyzed using anti-DUSP3 antibody (Abcam clone EPR5492), anti-DUSP1 antibody (Millipore 07-535), and anti-FLAG antibody (Santa Cruz Biotechnology sc-166355).

### Human PBMC isolation and transduction

Human whole blood from healthy anonymous volunteer donors was purchased from a NIH blood bank. This was exempted from the need for informed consent and Institutional Review Board review, as determined by the NIH Office of Human Subjects Research Protection. Human peripheral blood mononuclear cells (hPBMC) were isolated from whole blood by Ficoll density gradient separation, cultured in complete RPMI medium mentioned above, and stimulated with 1 μg/mL anti-CD3 (BioLegend 300334) and 1 μg/mL anti-CD28 (BioLegend 302944) soluble antibodies for 18 h. Purified and concentrated lentiviral particles were then added to the hPBMC culture, with 8 μg/mL Polybrene (Millipore Sigma TR-1003) and 100 U/mL recombinant human IL-2 (TECIN (teceleukin)). These cells were then spun at 400 X *g* for 90 minutes in a pre-warmed centrifuge at 34 °C.

## Results

### Dual-color Inducible CRISPR-Cas9 Editing System in RAW264.7 Macrophages

To achieve a controllable balance between on-target and off-target mutations induced by CRISPR, we adopted the doxycycline-inducible pCW-Cas9 system to enable precise manipulation of the genome editing time period. In pCW-Cas9-transduced RAW macrophages, a range of Cas9 protein expression was induced by doxycycline dose titration (Figure 1 - figure supplement 1A). We modified this existing system to enable single-cell flow cytometric analysis and developed the pCW-Cas9-2A-EGFP (iCE) lentiviral expression plasmid (Figure 1A). Doxycycline titration in the iCE-transduced and puromycin-selected RAW cells (RAWiCE) revealed saturating Cas9-2A-EGFP protein expression at 1.0 µg/mL doxycycline (Figure 1B). Single clone isolation from the RAWiCE line yielded cells that fully responded to doxycycline induction (Figure 1C), indicating the digital nature of this inducible expression system. To track single cells that constitutively express a targeting gRNA with a different set of antibiotic selection and live cell fluorescent markers, we developed the LentiGuide-TagBFP-2A-BSD (LGB) plasmid (Figure 1D) based on the LentiCRISPRv2 backbone, which had been optimized to produce high-titer lentiviral particles. For applications that require simultaneous tracking of two targeting gRNAs, we also developed the LentiGuide-TagRFP-2A-BSD (LGR) plasmid (Figure 1 - figure supplement 1B) to offer a different color option and enhance the overall flexibility of this inducible CRISPR-Cas9 editing system.

**Figure 1.**
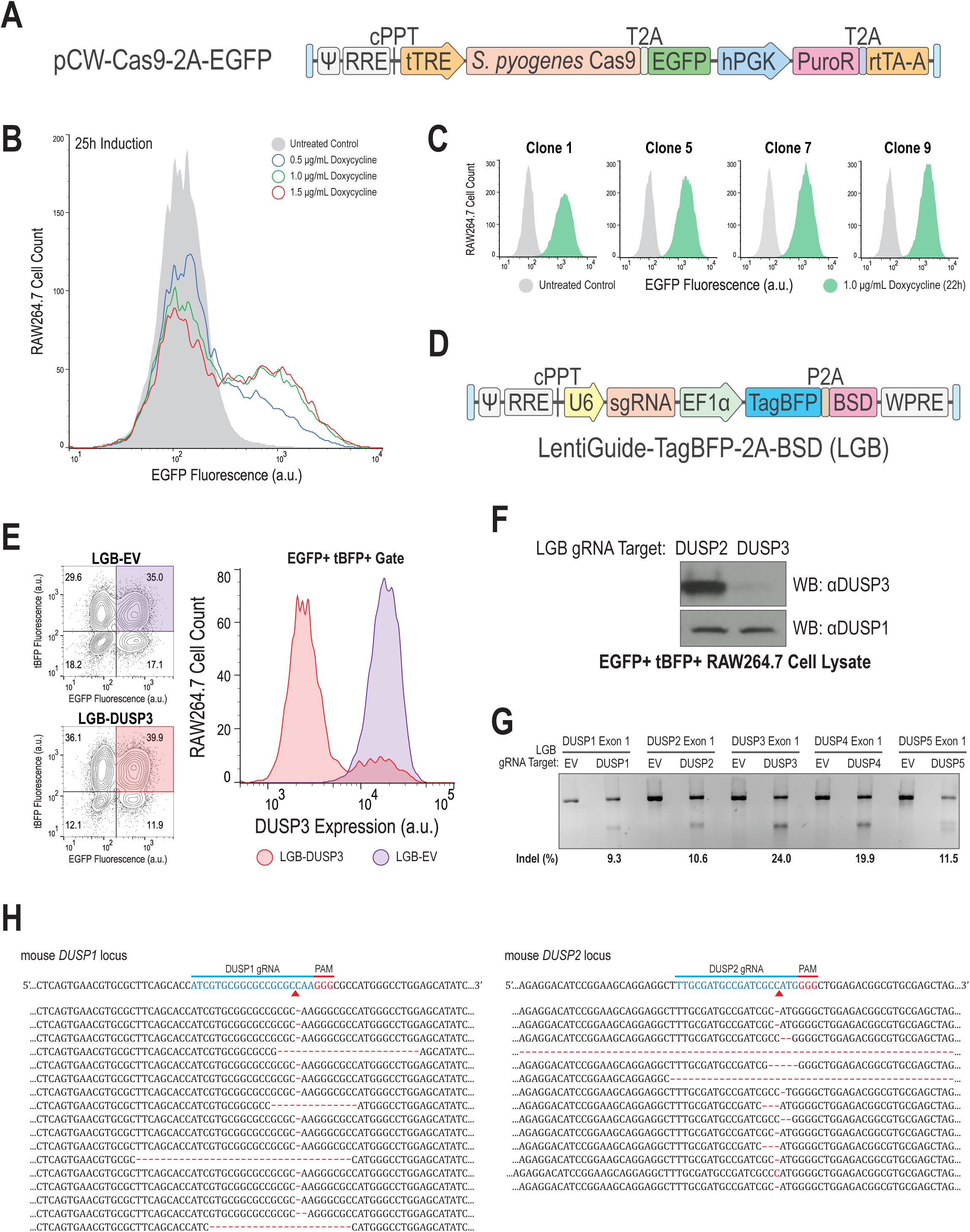
Dual-color inducible CRISPR strategy efficiently modifies DUSP genes in RAW264.7 macrophages. (A) Lentiviral vector encoding inducible Cas9-T2A-EGFP under the control of the tTRE promoter. (B) EGFP expression by RAWiCE cells treated with doxycycline. (C) EGFP expression by RAWiCE clones treated with doxycycline. (D) Lentiviral vector for gRNA expression with TagBFP fluorescent marker and blasticidin resistance gene BSD. (E) Analysis of DUSP3, EGFP and tBFP by RAWiCE cells transduced with the indicated LGB lentiviral particles and selected with blasticidin after doxycycline treatment. Results are representative of three independent experiments. (F) DUSP protein expression by RAWiCE cells transduced with the indicated LGB targeting DUSP2 or DUSP3. Data are representative of three independent experiments. (G) Indel analysis of genomic DNA from RAWiCE lines. (H) Sequence analysis of the genomic DNA from the RAWiCE lines.

To test this method, we targeted a family of dual-specificity phosphatases (DUSPs) in RAW cells. The RAWiCE cells transduced with LGB targeting each DUSP genomic locus were first selected by blasticidin to maximize the gRNA-expressing (tBFP+) population. Following doxycycline induction at 1.0 µg/mL for 6 days, we analyzed DUSP3 expression by intracellular staining and EGFP+ tBFP+ gating (Figure 1E). We observed substantial (>95%) reduction in total DUSP3 protein expression (Figure 1F) within the CRISPR-targeted EGFP+ tBFP+ population, indicating that the majority of targeted cell clones (Figure 1E) were indeed complete knockouts (KO) of DUSP3 expression. Due to limited antibody availability, we performed T7 Endonuclease I (T7EN) assays to confirm the presence of insertion/deletion (indel) mutations at the targeted genomic loci (Figure 1G). Since the T7EN assay does not yield the actual DUSP3 KO percentage, we performed genomic DNA sequencing to better estimate the CRISPR-editing outcomes of all five DUSP genes. The sequencing results revealed that most if not all targeted DUSP alleles had been mutated at the expected loci surrounding the PAM (protospacer adjacent motif) sites (Figure 1H & Figure 1 - figure supplement 1C). Overall, these results strongly suggest that our dual-color inducible CRISPR strategy confers high genome editing efficiency in an extremely sensitive, albeit transformed, innate immune cell type.

Upon stimulating DUSP-KO RAW cells with TLR4 ligand, we did not observe sustained p38 phosphorylation in DUSP1-KO RAW cells (Figure 2 - figure supplement 1A), an expected phenotype based on a previous study using primary BMDM derived from DUSP1-KO mice (38). Thus, we examined the effects of such a KO in BMDM using the same gRNA used to target DUSP1 in RAW cells. Although lentiviruses can transduce non-dividing cells, transduction efficiency is optimized in cells undergoing active replication. Based on the Ki67 proliferative marker expression profile of bone marrow cells cultured with M-CSF (Figure 2 - figure supplement 1B), we developed a lentiviral gRNA transduction protocol (Figure 2A) to create target gene KO BMDM for further analysis. Unlike RAW cells, BMDM are more sensitive to crude lentiviral supernatant from HEK293T packaging cells seen as a dramatic post-transduction decline in the live BMDM percentage (Figure 2B). To optimize BMDM survival, we further purified and concentrated lentiviral particles to remove potentially toxic materials from the crude supernatant. This approach improved the live BMDM percentage as the LGB titration reached a peak transduction efficiency (Figure 2B). Using this improved transduction method for LGB expression in Cas9+ EGFP+ BMDM, we performed intracellular protein staining and validated the editing efficiency of DUSP3 gRNA (Figure 2C). Consistent with previous report using BMDM from DUSP1-KO mice (38), we observed sustained TLR4-induced p38 phosphorylation (p-p38) in BMDM transduced with the LGB-DUSP1 gRNA expression, as compared to the EV (empty vector) and the DUSP4 controls (Figure 2D). Increased p-p38 phenotype generally correlated with the LGB-DUSP1 gRNA expression level, as indicated by tBFP fluorescence in Cas9+ EGFP+ BMDM (Figure 2E). Our results suggest that TLR4-induced p38 is differentially regulated between primary BMDM and the transformed RAW macrophages and illustrate the utility of our method for study of these otherwise hard-to-modify cells.

**Figure 2.**
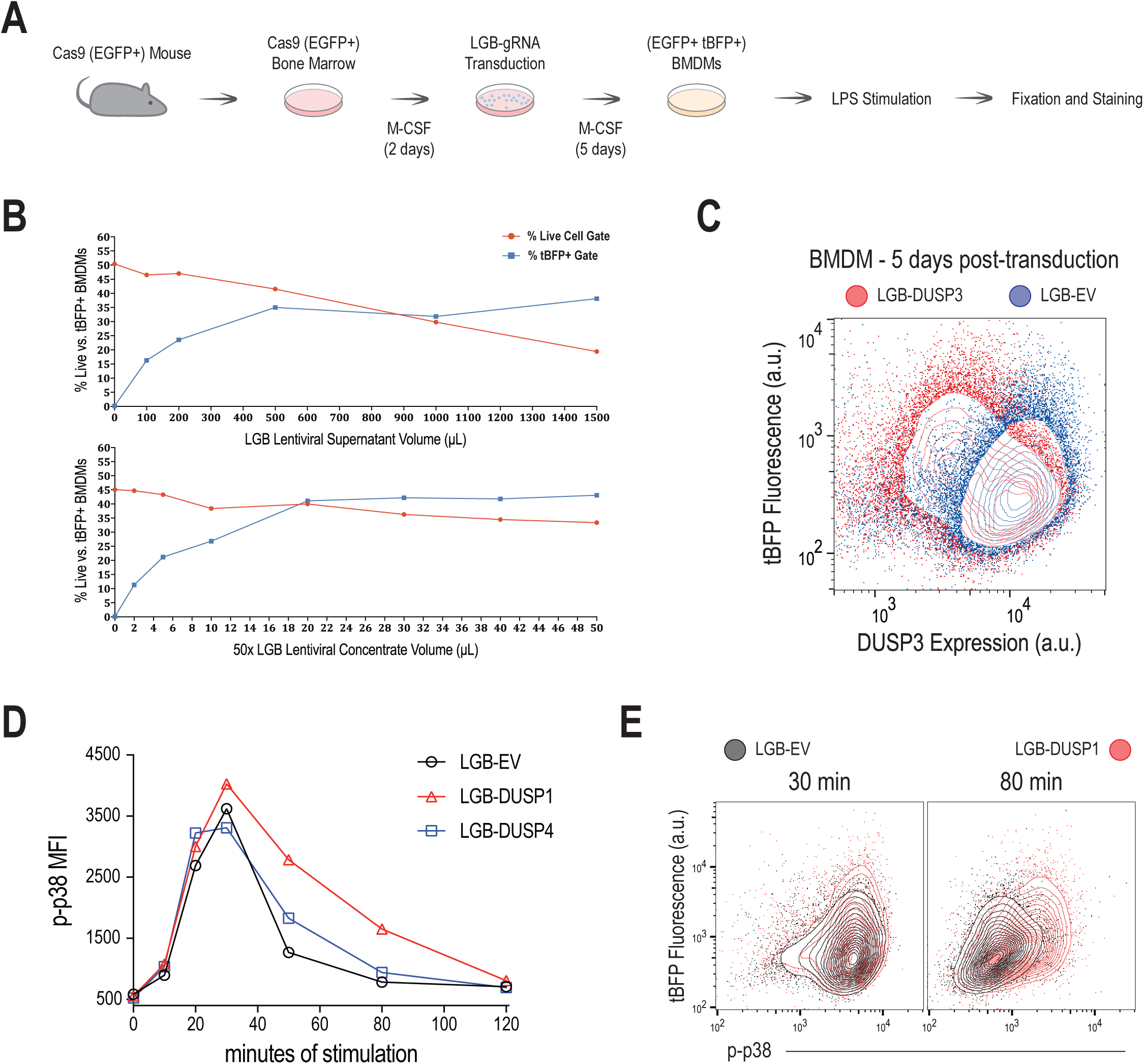
LGB-mediated CRISPR in BMDM recapitulates reported role for DUSP1 in control of TLR4-p38 signaling. (A) Workflow diagram for generating CRISPR-KO BMDMs for TLR4 signaling pathway analysis. (B) Lentiviral titration in BMDMs for cell toxicity. (C) DUSP3 protein expression following LGB-DUSP3-transduction of Cas9+ BMDMs. (D) Phospho-p38 induced by TLR4 engagement of DUSP1-KO BMDMs as compared to the EV and the DUSP4-KO controls. Data are representative of two independent experiments. (E) Examination of TLR4-induced phospho-p38 signaling together with LGB-DUSP1 gRNA expression as indicated by tBFP fluorescence.

### An Efficient Selection Strategy for CRISPR-induced HDR Clones in Jurkat T Cells

We next turned to new strategies for editing the genomes of lymphocytes. To minimize off-target effects, we designed a pair of gRNAs for targeting with the Cas9-D10A nickase (Cas9n) strategy, choosing the human STAT3 locus as an exemplar (Figure 3A). This pair of gRNAs together with Cas9n expression in HEK293T cells was confirmed to create indel mutations at the targeted STAT3 loci, as measured by the T7EN assay (Figure 3B) following co-transfection of the PX461 and the pKLV-BFP plasmids. However, our goal was more ambitious than just achieving gene expression loss in these cells. In addition to expression of Cas9n and a pair of gRNAs lying in close proximity, a DNA donor template is required to induce precise knock-in modification via the HDR repair mechanism. The STAT3 HDR donor plasmid (Figure 3C) was designed with 2000-bp homology arms flanking the insert fragment containing the puromycin resistance (PuroR) gene, followed by the T2A sequence and the mNeonGreen (mNG) fluorescent protein-coding gene. Following co-transfection of PX461-STAT3-gRNA-A, pKLV-BFP-STAT3-gRNA-1 and STAT3 HDR donor plasmids, approximately 6e5 total (<0.5% co-transfection efficiency) Jurkat T cells were selected with puromycin and separated by dilution cloning before further validation (Figure 3D). Based on the low HDR efficiency for large fragment insertion (<10%), it is technically challenging to isolate this tiny (<0.05% or <300 cells) population of successful HDR cell clones without using any selection method. After puromycin selection, flow cytometric analysis of 40 puromycin-resistant Jurkat cell clones revealed that most of the selected population had acquired mNG expression to various extents (Figure 3E). Further genomic PCR analysis of 7 clones that express medium to high level of mNG showed that 6 out these 7 clones were indeed complete knock-in for all STAT3 alleles (Figure 3F). Based on these results, we estimated the clonal distribution of the parental mNG-STAT3 Jurkat cell line (Figure 3G). The high percentage (82.5%) of partial or complete mNG-STAT3 knock-in clones suggests that this kind of antibiotic selection strategy efficiently enriches the desired HDR cell clones.

**Figure 3.**
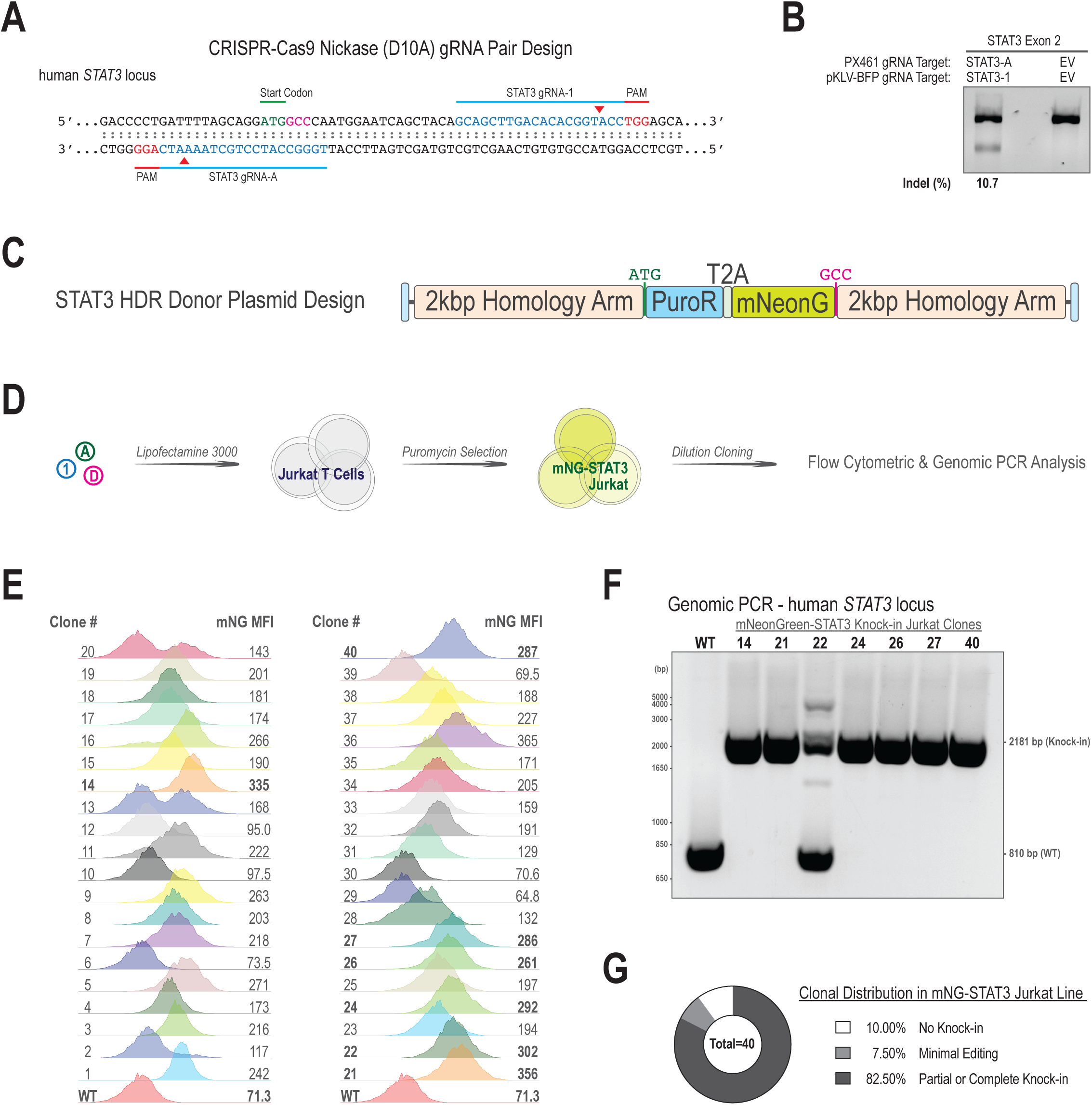
In-frame PuroR HDR donor design allows highly specific selection of mNG-STAT3 knock-in Jurkat T cells. (A) Cas9 nickase gRNA pair design targeting human STAT3 locus to induce HDR events near the start codon. (B) T7EN assay for locus-specific editing in HEK293T cells. (C) STAT3 HDR donor plasmid. Start codon and the original second codon are labeled as shown in A. (D) Strategy for puromycin selection of clonal Jurkat cells with the STAT3 locus-targeted Jurkat T cells. (E) mNeonGreen expression by clones of puromycin-resistant mNG-STAT3 Jurkat cells. (F) PCR analysis of the hSTAT3 locus in targeted mNG-STAT3 Jurkat cells. (G) Targeting efficiency in the mNG-STAT3 line.

We extended studies of this method by targeting the human BCL10 locus to express a fluorescent protein-tagged fusion protein, using 6 nickase gRNA pairs (Figure 4A) and measuring the targeting efficiency by T7EN assay (Figure 4B) as in Figure 3B. Despite the higher indel frequency induced by the D3 gRNA combination (Figure 4A & 4B), we selected the highest-scoring (crispr.mit.edu) A1 gRNA combination to minimize potential off-target effects. To enhance co-transfection efficiency, we created a double gRNA expression plasmid (Figure 4C) based on the PX462 backbone. Furthermore, we attempted to minimize the genome editing footprint by incorporating loxP sites flanking the PuroR-T2A selection cassette in the BCL10 HDR donor plasmid (Figure 4D). Upon CRE-mediated excision of the loxP-PuroR-T2A-loxP fragment, further genome editing with the same strategy can be applied because puromycin sensitivity is restored (Figure 4E). After plasmid co-transfection and puromycin selection, most Jurkat cells had acquired mNG-BCL10 expression (Figure 4F) and showed dispersed protein localization, as compared to the nuclear H2B-BFP marker (data not shown). After NIL-CRE (non-integrating lentiviral particles; see Figure 5A for details) treatment, we observed enhanced mNG-BCL10 expression level (Figure 4G) and a substantially higher frequency of mNG-BCL10 aggregate formation (39) upon antigen receptor stimulation (data not shown). Puromycin re-treatment killed approximately 99.8% of the NIL-CRE-treated mNG-BCL10 Jurkat cells but did not affect the survival of the original mNG-BCL10 line (data not shown). Finally, based on this NIL-CRE-treated or Puro-T2A-excised mNG-BCL10 Jurkat parental cell line, we repeated STAT3 genome editing as in Figure 3 but using a modified STAT3 HDR donor plasmid (Figure 4H) in which the mScarlet element replaces the mNeonGreen element (Figure 3C). Following plasmid co-transfection and puromycin selection, more than 75% of mNG-BCL10+ Jurkat cells had gained the targeted mScarlet-STAT3 expression (Figure 4I). With minimal off-target editing concern, these results demonstrated a highly efficient yet sustainable selection method to precisely enrich targeted knock-in cell population from very few recombinant clones created by CRISPR-mediated HDR.

**Figure 4.**
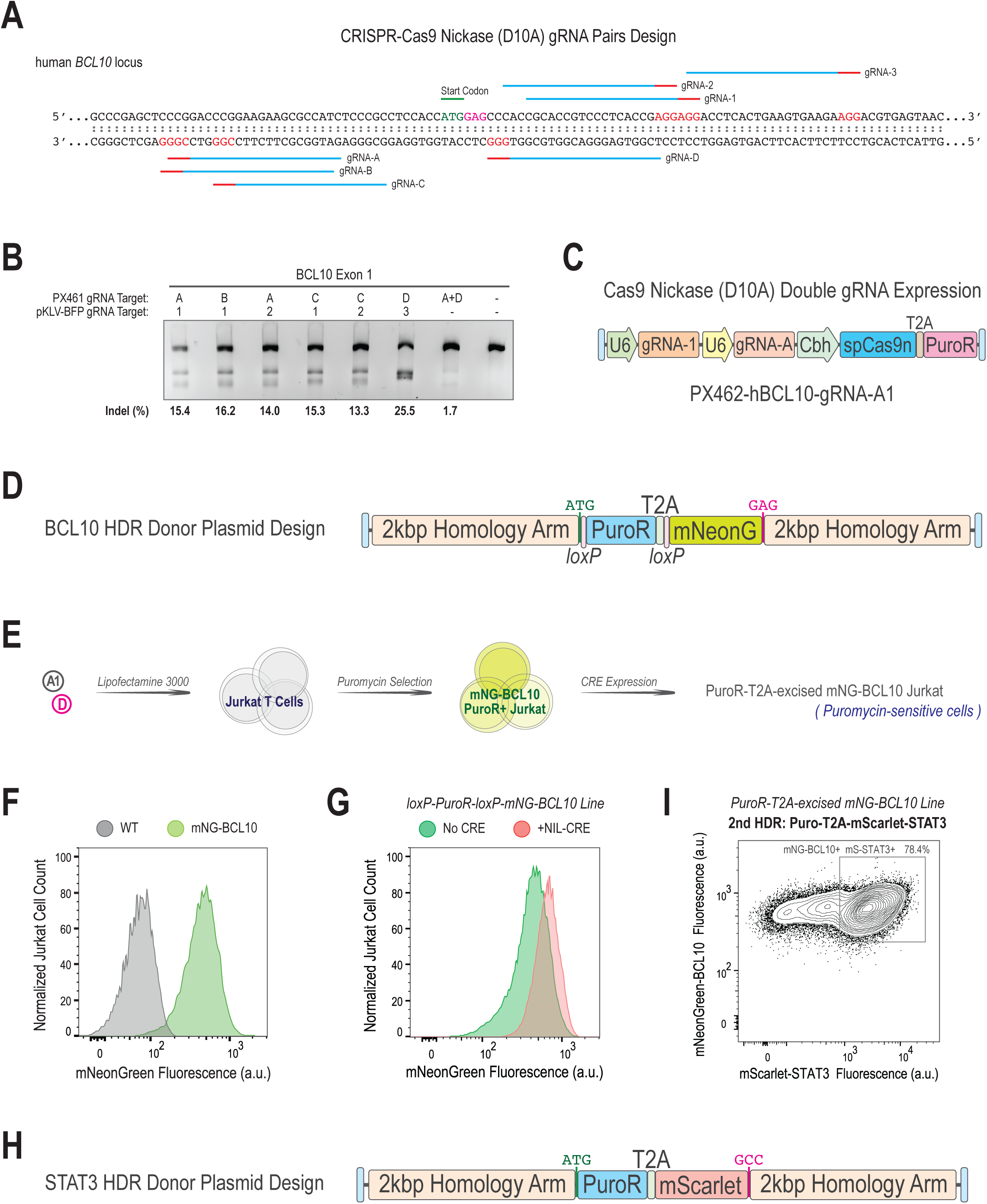
CRE-loxP excision strategy minimizes genomic footprint of the PuroR selection cassette and allows for multiple rounds of genome editing. (A) Cas9 nickase gRNA pair design targeting the human BCL10 locus. (B) T7EN assay for locus-specific editing in HEK293T cells. (C) Double gRNAs and Cas9-D10A nickase expression plasmid derived from the PX462 backbone. (D) BCL10 HDR donor plasmid design. (E) Strategy for creating puromycin-selected Jurkat T cells with post-selection excision of the Puro-T2A cassette. (F)(G) mNeonGreen expression by puromycin-selected mNG-BCL10 knock-in Jurkat line with or without NIL-CRE treatment. (H) Modified STAT3 HDR donor plasmid. (I) mNeonGreen and mScarlet co-expression by puromycin-selected mScarlet-STAT3/mNG-BCL10 double knock-in Jurkat line.

### All-in-one Non-integrating Lentiviral Delivery of the CRISPR-HDR Blueprint

As an alternative approach, we sought to deliver Cas9, gRNA, and the HDR donor template to the entire T cell culture in a transient transfection manner. We therefore turned to the use of integrase-deficient lentiviral particles. We packaged the Lenti-CMV-mCherry-P2A-CRE viral vector either with the wild-type integrase in the psPAX2 plasmid to produce integrating lentiviral (IL) particles, or with the integrase-deficient mutant in the psPAX2-D64V plasmid to produce non-integrating lentiviral (NIL) particles. While the IL transduction created stable mCherry expression in Jurkat T cells, the NIL delivery failed to integrate into the genomes and led to transient mCherry expression (Figure 5A). To test whether NIL delivery of the entire CRISPR-HDR instruction can generate targeted knock-in cells, we assembled the LentiCRISPRv2B-HDR-mCherry-2A-hCD3E viral vector (Figure 5B) and examined two gRNAs targeting the human CD3E locus (Figure 5C). 7 days after the NIL-CRISPR-Cas9-HDR delivery, we indeed observed a distinct population of mCherry+ Jurkat cells by flow cytometry (Figure 5D). Using primers that distinguish the endogenous locus from randomly-integrated HDR template sequences, genomic PCR analysis confirmed the presence of targeted knock-in alleles within the mCherry-sorted Jurkat population (Figure 5E).

**Figure 5.**
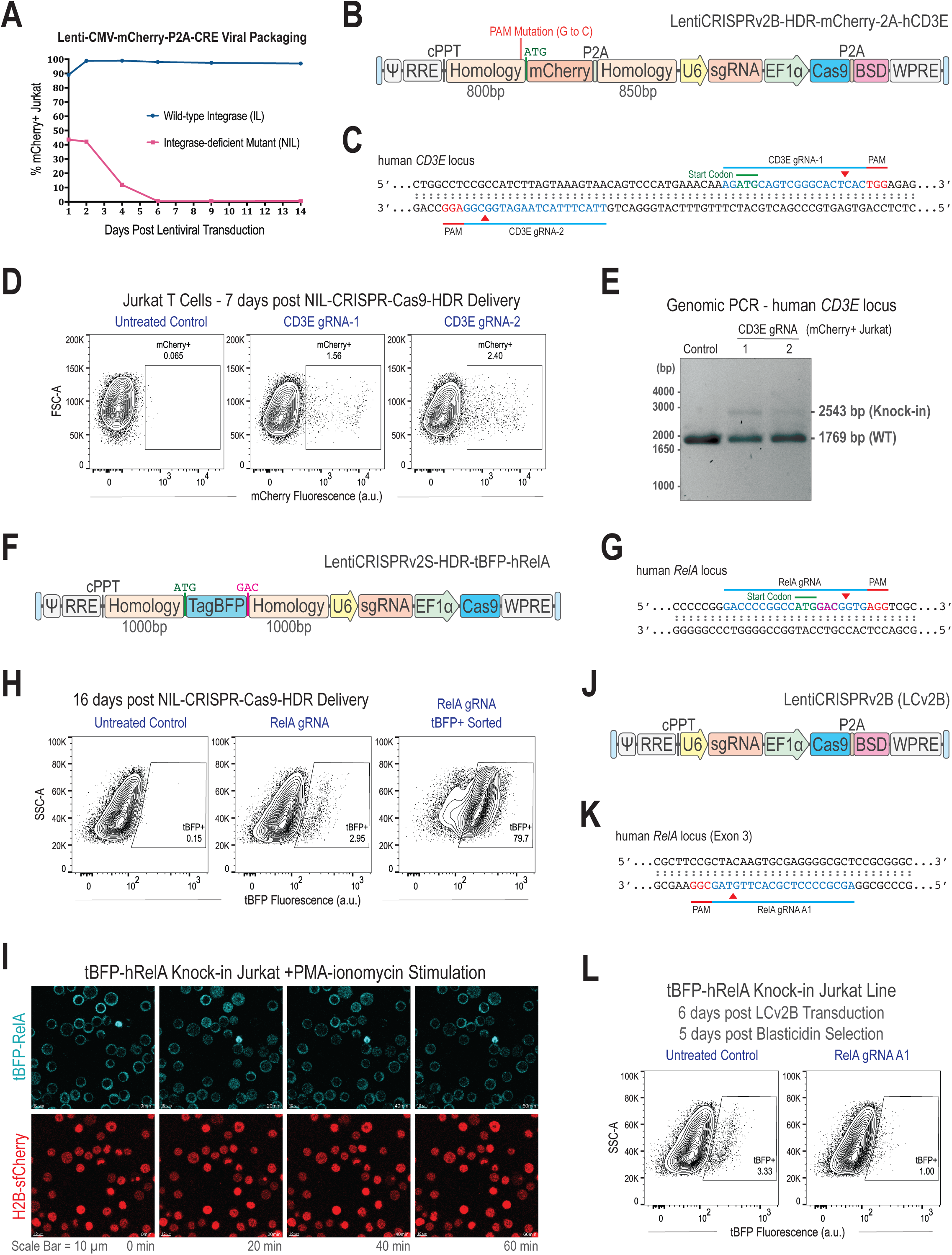
All-in-one NIL-CRISPR-Cas9-HDR delivery enables targeted genome modifications in human Jurkat T cells. (A) Time course of mCherry-P2A-CRE expression after NIL vs. IL transduction of Jurkat T cells. (B) All-in-one lentiviral design for mCherry-P2A-CD3E knock-in via CRISPR-Cas9-HDR. PAM site for CD3E gRNA-2 was modified from AGG to ACG in the left homology arm. (C) Human CD3E locus showing targeting sites of both gRNAs tested. (D) mCherry expression by Jurkat cells transduced with NIL-LCv2B-HDR-mCherry-2A-hCD3E. Results are representative of two independent experiments. (E) PCR analysis of homology arm regions of mCherry+ Jurkat cells. (F) All-in-one lentiviral design for tBFP-RelA knock-in via CRISPR-Cas9-HDR. Start codon and the original second codon are labeled as highlighted in G. (G) Human RelA locus showing the target site of gRNA used in F. (H) tBFP expression of Jurkat cells untreated or treated with LCv2S-HDR-tBFP-hRelA, before or after tBFP+ cell sorting. Results are representative of two independent experiments. (I) Live cell confocal time-lapse imaging of tBFP-RelA knock-in Jurkat cells stimulated with PMA-ionomycin. Pre-existing H2B-sfCherry expression created by lentiviral transduction was used as nuclear marker. (J) Modified lentiCRISPRv2 plasmid LentiCRISPRv2B replacing puromycin resistance with blasticidin resistance gene BSD. (K) Human RelA locus showing the target site of gRNA A1 near exon 3. (L) tBFP expression by the Jurkat tBFP-RelA knock-in line either untreated or transduced with LCv2B carrying RelA gRNA-A1 to knockout endogenous RelA expression.

Using NIL delivery of the LentiCRISPRv2S-HDR-tBFP-hRelA viral vector (Figure 5F), we attempted to generate a TagBFP-RelA fusion knock-in Jurkat cells, employing a gRNA targeting the human RelA locus (Figure 5G). This HDR attempt yielded a distinct and stable tBFP+ Jurkat population that was further purified by FACS (Figure 5H). We used live cell confocal time-lapse imaging to visualize the RelA nuclear translocation response to PMA-ionomycin stimulation (Figure 5I). As expected, in the steady-state tBFP-RelA was mostly excluded from the H2B-sfCherry-defined nuclear region. Upon NF-κB pathway activation, a rapid but transient increase in the nuclear to cytoplasmic ratio of tBFP-RelA was observed in approximately 85% of the Jurkat population (Figure 5I). To distinguish targeted genomic knock-in from random lentiviral integration, we transduced the original tBFP-RelA knock-in Jurkat line with LentiCRISPRv2B (Figure 5J) carrying RelA gRNA-A1 (Figure 5K) to inactivate endogenous RelA expression. After 5 days of blasticidin selection, approximately 70% of the gated population had lost tBFP expression (Figure 5L), indicating that most of the tBFP+ cells were indeed created by the HDR mechanism at the human RelA locus. Taken together, these results provide proof-of-concept data to support the NIL delivery approach as a powerful alternative to the existing CRISPR-HDR methods in human T cells.

### Selective Gene Expression Induction with All-in-one Lentiviral CRISPRa System

The vectors and methods described above are useful for either eliminating expression of a particular gene or modifying an already-expressed gene in a given cell type, but there are numerous situations in which it would be useful to induce expression of an otherwise silent locus in a specific cell type. To achieve this end, we created an all-in-one lentiviral CRISPRa system based on an existing two-plasmid system (32) and we refer it as LentiSAMPHv2 (Supplementary Figure A). With this system, we tested 6 gRNAs from an existing CRISPRa library (32) targeting the promoter region of the human PDCD1 or PDL1 genomic locus. These gRNAs induced various levels of target gene expression (Supplementary Figure B), as compared to the corresponding no gRNA control condition. For the gRNA that induces the highest level of PD1 expression, we also tested the specificity of this induction by measuring the cell surface expression level of PDL1 (Supplementary Figure C).

## Discussion

Many research groups have encountered failures in their unpublished CRISPR-KO attempts to engineer the widely-studied RAW macrophage cell line (personal communications). Using the lentiCRISPR system, we consistently observed massive cell death following lentiviral transduction and puromycin selection, likely due to a combinatorial effect of low viral titer and cytotoxic materials from the crude lentiviral supernatant. In contrast, the high transduction efficiency of our LGB gRNA expression system ensures that a majority of the transduced RAW cells survive under blasticidin selection, conferring a good representation of the original cell line. Inducible Cas9 expression allows more precise control of the genome editing time window and therefore minimizes long-term off-target effects that cannot be avoided by constitutive Cas9 expression in many lentiviral-based CRISPR expression systems. For gRNAs that efficiently knockout essential cell survival genes or oncogenes, percentage changes in EGFP+ tBFP+ cells over doxycycline treatment time help quantify the impacts of these gRNAs.

The high genome editing efficiency and the rapid single-cell phenotyping option of our dual-color inducible CRISPR system in RAW cells makes this platform suitable for high-throughput genome-wide gRNA library screening. Our inducible CRISPR system allows for stable uninduced cell lines to be frozen for long-term storage and future analysis. Furthermore, we have optimized these methods for use in primary immune cells. Lentiviral or retroviral supernatant produced from HEK293T packaging cells often influences target cell growth rate and morphology, especially when a high volume-ratio of viral supernatant to cell culture medium is applied to the target cell line. Our data illustrate that further purification of lentiviral particles helps minimize BMDM cell death. Thus, our methods achieve high transduction efficiency without sacrificing cell health in culture. With these approaches, we were able to identify Cas9 and gRNA double positive macrophages, using both the RAW murine cell line and primary BMDM. Interestingly, these studies revealed distinct MAPK regulation between these cells. CRISPR deletion of the phosphatase DUSP1 in BMDM resulted in sustained p38 phosphorylation, consistent with published studies in BMDM generated from DUSP1 deficient mice (38). In contrast, DUSP1 deletion in RAW cells, using the same gRNA, did not yield detectable changes in p38 activation. This adds to our previous reports of dysregulated TLR4-induced MAPK activity in RAW cells compared to BMDM (40).

The Cas9 nickase strategy requires two gRNA targets to be in close proximity to efficiently induce DNA double-strand break repair mechanisms (8,9). Therefore, it is extremely unlikely to find another off-target pairing site in the genome. Our data support previous reports that the pair of Cas9 nickases are optimized with minimized offset of the gRNA pairs (8,9). Without including an upstream promoter in the donor template design, our PuroR selection strategy reduces the probability of selecting randomly integrated clones. Obviously, this strategy relies on genomic loci that are well expressed to generate sufficient expression of the drug resistance gene. Among the target genes that we have tested, we observed that this strategy worked best with the N-terminal PuroR-2A but not the C-terminal 2A-PuroR fusion expression. Combined with a CRE-lox or FLP-FRT recombination design, our PuroR selection strategy is further optimized to sustain more rounds of genome editing, enabling sequential engineering of multiplex reporters, significantly reducing the genomic footprint to a short single recombination site. Following the NIL-mediated mCherry-P2A-CRE or mScarlet-P2A-CRE (Figure 5 - figure supplement 1D), or commercially available Cre Gesicles (Takara) delivery, target cells that transiently express CRE recombinase can be isolated from the untreated population via fluorescence-activated cell sorting (FACS). For relatively small proteins such as BCL10, the protein expression level can be dramatically reduced by the genomic insertion of the PuroR selection cassette together with a fluorescent protein. Our data suggest that reduced BCL10 protein expression can lead to the loss of BCL10 oligomerization upon antigen receptor signaling, which justifies the need to remove the PuroR selection cassette to achieve a more functional BCL10 reporter Jurkat T cell line.

Lentiviruses such as HIV have evolved to efficiently infect human T cells, making them perfect delivery vehicles for human T cell genome engineering. In Jurkat cells, lentiviral delivery is much more efficient than any existing transfection method including electroporation, which often leads to substantial cell death upon electric shock and reduces cell growth and overall protein production for the surviving population. In addition, lentiviral particles are easy to prepare by most laboratories, carry double to triple the payload as compared to the AAV system, and therefore enable all-in-one design and delivery of a Cas9, gRNA expression and HDR donor template. Although there is now an existing report (41) that demonstrates the feasibility of NIL delivery of the CRISPR-HDR blueprint, we now demonstrate the utility of this method in an especially difficult to engineer type of cell, T lymphocytes. Our experiments using human primary T cells showed that the NIL-LCv2R (Figure 5 - figure supplement 1B) particles efficiently transduced more than 80% of the total population (Figure 5 - figure supplement 1E).

As one of the first studies to report the use of NIL as delivery vehicle for CRISPR-HDR applications, we suggest many exciting optimization possibilities for this method. First of all, sensitive marker expression such as mScarlet in the LCv2R backbone (Figure 5 - figure supplement 1B) or tEGFR in the LCv2E backbone (Figure 5 - figure supplement 1C) could be included to eliminate residual random integration events from NIL particles. In addition, various HDR-stimulating chemicals such as RS-1 (27) could be tested to enhance knock-in efficiency. Furthermore, eSpCas9 or more specific Cas9 variant could be used instead of Cas9 nuclease to minimize off-target effects. Finally, the use of both NIL and AAV systems could be combined to deliver few copies of NIL-CRISPR-Cas9-gRNA but high copy number of AAV-HDR donor template (28) per target cell to maximize HDR efficiency and donor template accuracy.

We also introduce here a new all-in-one vector system for activation of selected gene loci in immune cells. Using a single lentivirus containing gRNA that targets the human PDCD1 or PDL1 genomic locus, we demonstrate high level expression of a previously silent locus in Jurkat cells and the value of this induced expression in assays of immune function. This strategy builds on pre-existing methods for large scale gene induction by providing experimentalists with a simple method for selective, one-molecule-at-a-time alteration of cell phenotype in a positive rather than negative manner.

Among the published reports using various CRISPR technologies in immune cells, there has been a lack of focus on using more accurate genome editing tools. As more precise CRISPR-based genome editing tools are being developed and characterized, future immune cell genome engineering studies should pay more attention to minimizing off-target editing effects. This is particularly important for immune cells that are modified for therapeutic applications that require the highest safety level of genome engineering.

## Acknowledgements

We thank Clinton Bradfield, Iain Fraser, and Wanjing Shang for critical reading of the manuscript. We thank Cindi Pfannkoch, Bin Lin, Caleb Ng, Sebastian Montalvo, Iain Fraser, and Michael Lenardo for helpful discussions and advice. This research was supported by the Intramural Research Program of the NIH, NIAID.

## Competing interests

All authors declare no competing interests.

---

**Figure 1 - figure supplement 1.**
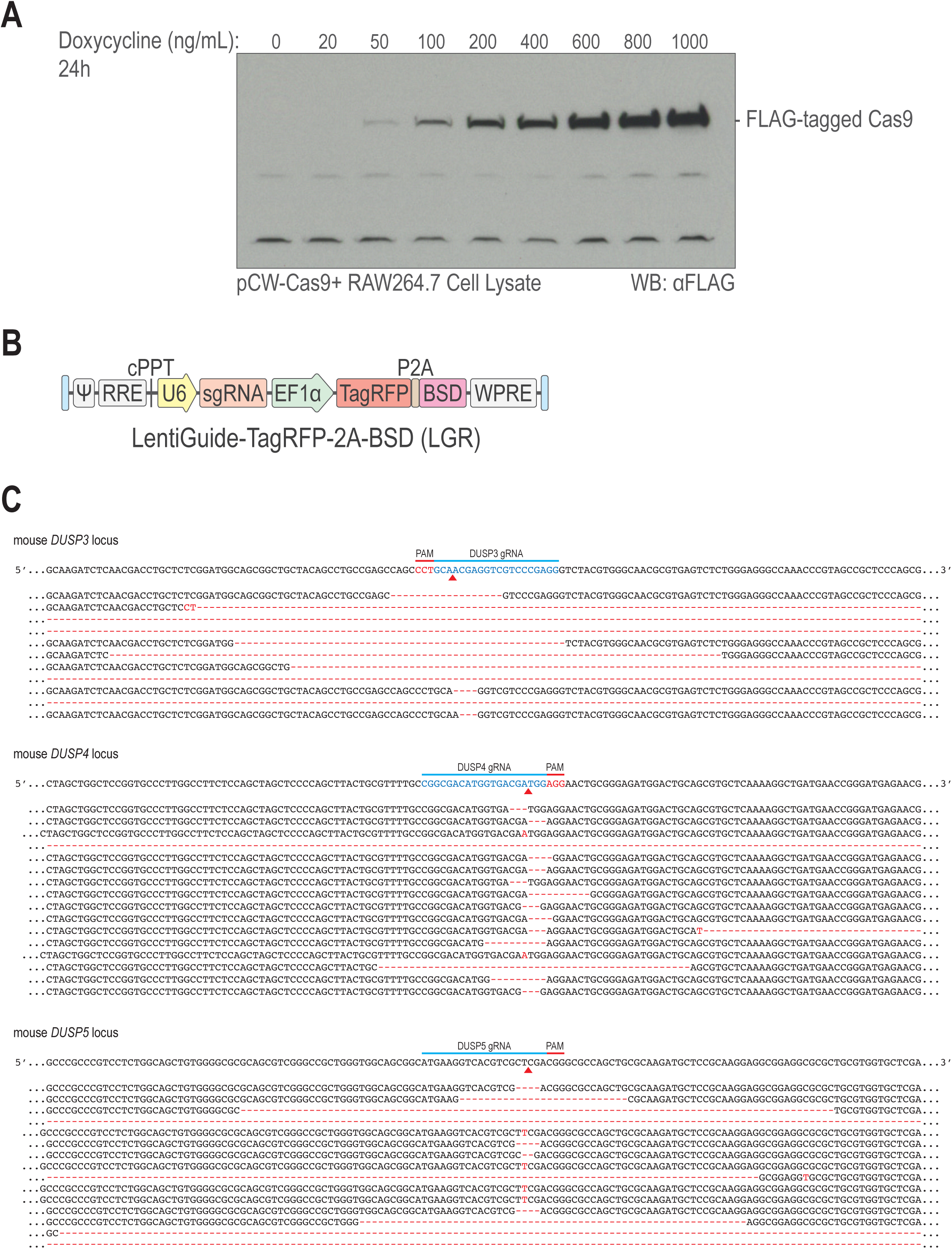
Inducible genome editing system in RAW264.7 macrophages. (A) RAW cells transduced with pCW-Cas9 were treated with titrating doses of doxycycline. Cell lysates were analyzed by western blot with anti-FLAG antibody. (B) Lentiviral vector for gRNA expression with TagRFP-T fluorescent marker and blasticidin resistance gene BSD. (C) Sequence analysis of the genomic DNA from the RAWiCE lines.

**Figure 2 - figure supplement 1.**
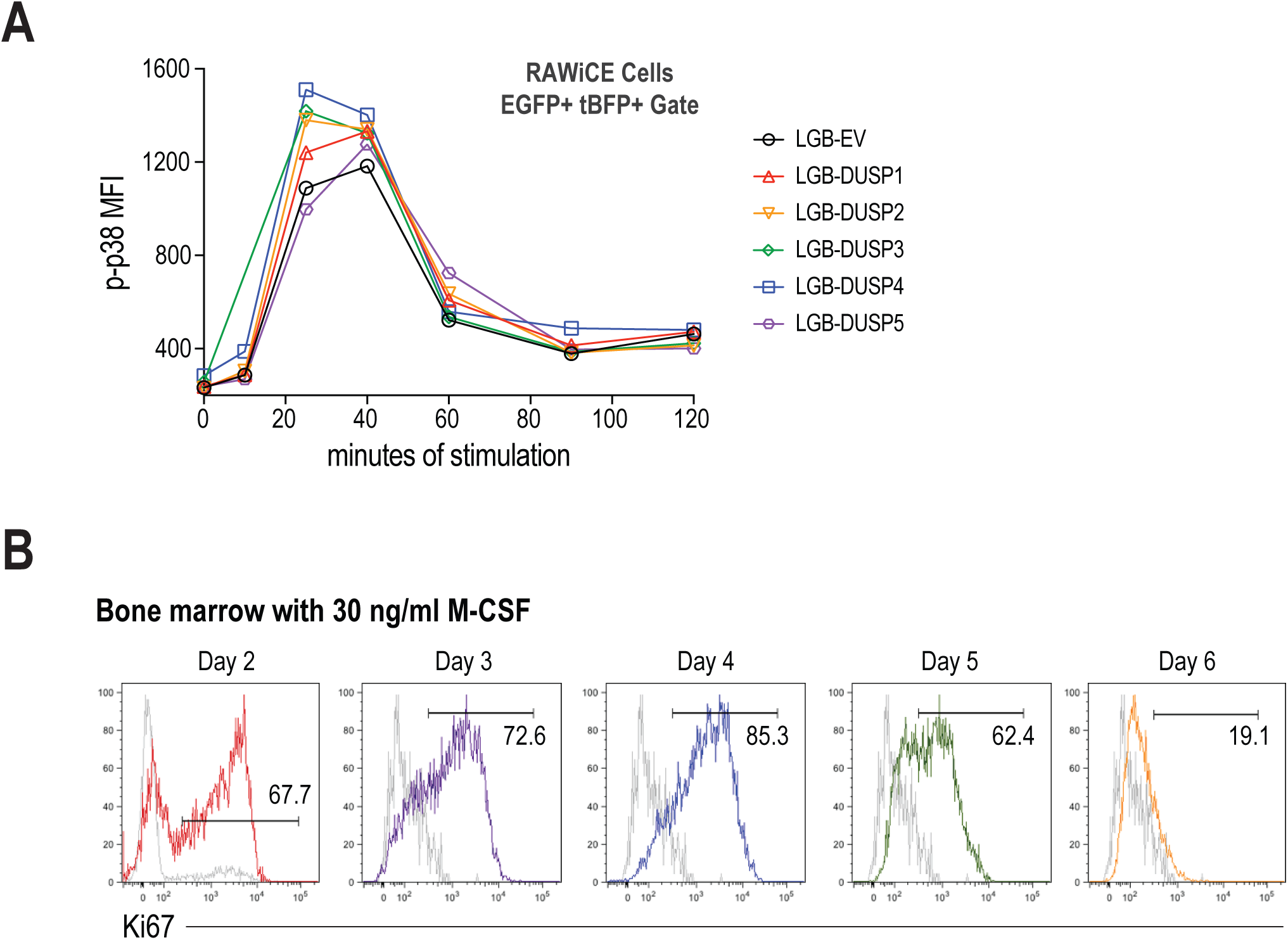
DUSP1-KO RAW cell phenotype and Ki67 profile of B6 bone marrow. (A) After blasticidin selection and doxycycline induction, RAWiCE cells transduced with the indicated LGB variant were stimulated with TLR4 ligand and mean phospho-p38 was quantified over time. Data are representative of two independent experiments. (B) Ki67 staining profile of C57BL/6 mouse bone marrow cells after 30 ng/mL M-CSF treatment.

**Figure 5 - figure supplement 1.**
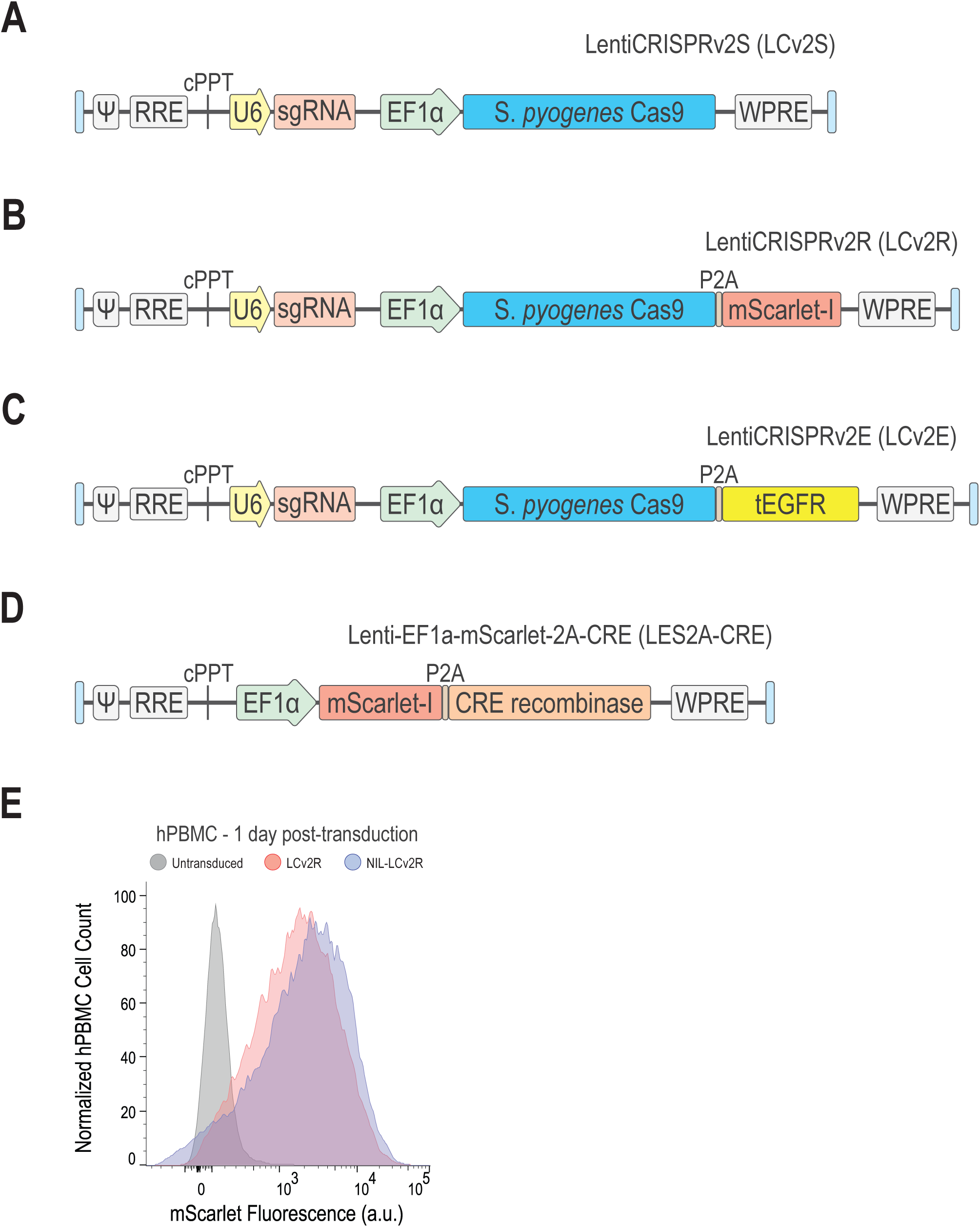
LES2A-CRE, LentiCRISPRv2 variants LCv2S, LCv2R and LCv2E, and lentiviral transduction efficiency in human PBMC. (A) The LentiCRISPRv2S plasmid offers reduced lentiviral vector size and flexible cloning of the marker of interest. (B) The LentiCRISPRv2R plasmid contains mScarlet-I for sensitive detection of Cas9 expression. (C) The LentiCRISPRv2E plasmid contains the tEGFR cell surface marker to report Cas9 expression. (D) The Lenti-EF1a-mScarlet-2A-CRE plasmid allows sensitive reporting of CRE recombinase expression via the mScarlet-I fluorescent marker. (E) mScarlet expression by human PBMC transduced with LCv2R or NIL-LCv2R.

**Supplementary Figure.**
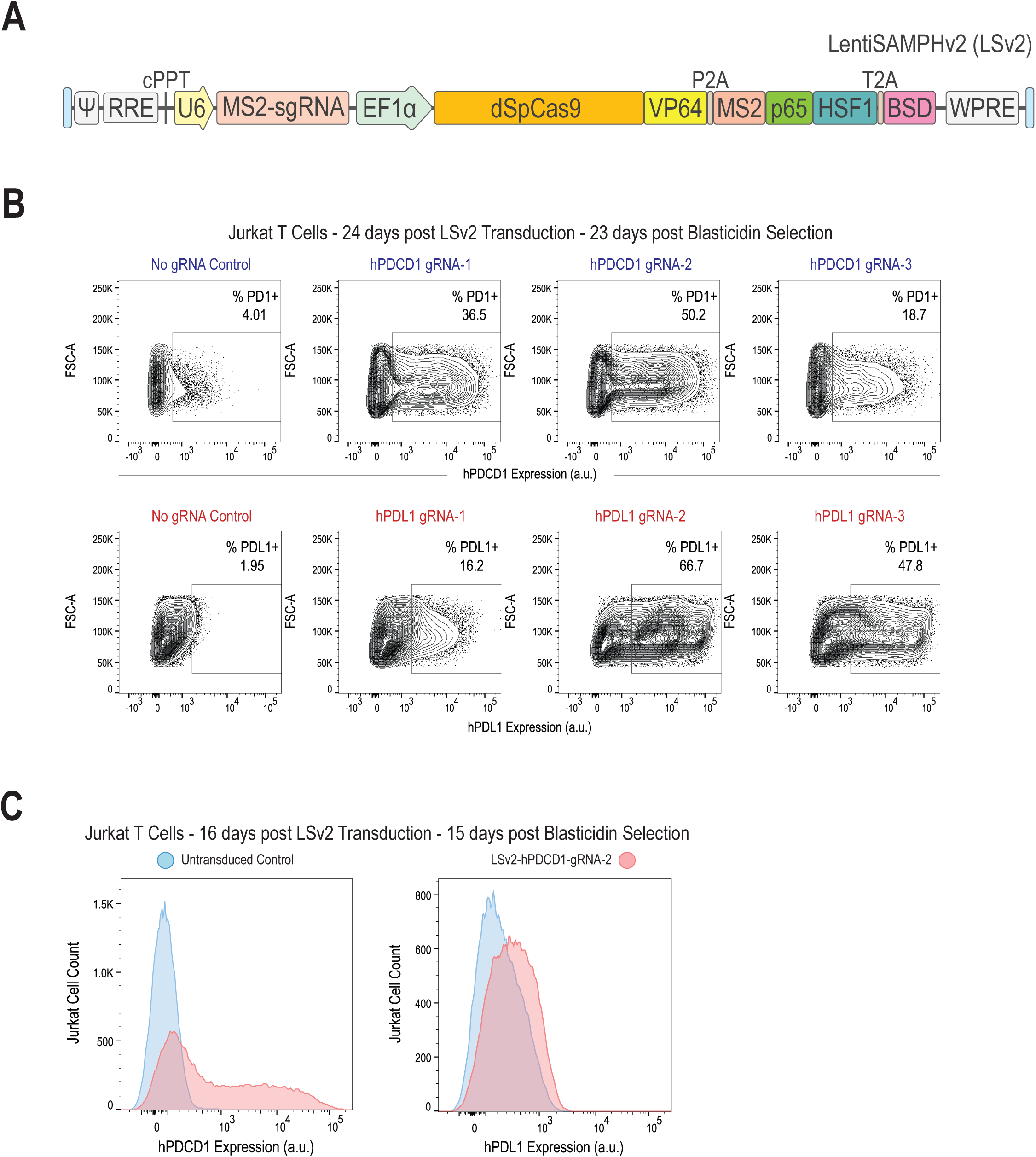
Lentiviral-mediated CRISPRa with the LentiSAMPHv2 System. (A) The all-in-one LentiSAMPHv2 (LSv2) plasmid for CRISPR-based gene activation applications. (B) Flow cytometric analysis of human PDCD1 (top panel) or PDL1 (bottom panel) cell surface protein expression after LSv2 transduction with empty vector control or the indicated CRISPRa targeting gRNA. (C) Flow cytometric analysis of human PDCD1 (left panel) or PDL1 (right panel) cell surface protein expression in untransduced control cells versus cells transduced with LSv2 targeting the human PDCD1 promoter region.

**Video 1. Live cell confocal time-lapse imaging data of tBFP-RelA translocations following PMA-ionomycin stimulation in human Jurkat T cells**.

## Notes

This work was supported by the Intramural Research Program of NIAID, NIH.

